# Temperature-modulated maternal effects vary with offspring developmental stage in *Drosophila melanogaster*

**DOI:** 10.1101/2024.04.26.591356

**Authors:** P Kohlmeier, B Van Schaik, I Pen, JC Billeter, TGG Groothuis

**Affiliations:** Groningen Institute for Evolutionary Life Sciences, University of Groningen, 9474AG Groningen, The Netherlands

**Keywords:** maternal effects, silver spoon effects, trans-generational plasticity, phenotypic plasticity, temperature, split brood design, Bayesian analysis, *Drosophila melanogaster*

## Abstract

An organism’s phenotype has the potential to vary in response to environmental factors, allowing it to adjust to environmental fluctuations. Maternal effects on offspring phenotypes have been recognized as important contributors to this phenotypic plasticity, although the extent and duration of this contribution remain elusive. At more advanced developmental stages, offspring may be able to assess their current environment more accurately than at earlier developmental stages, and therefore their reliance on maternal effects may decline over time. This study investigates how the magnitude and direction of maternal effects change between early and late development using the fruit fly *Drosophila melanogaster* as a model. We employed a split brood design to disentangle the effects of maternal ambient temperature from the effects of offspring ambient temperature at the larval vs. the pupal or early adulthood stage. We subsequently measured offspring phenotypes such as heat shock and cold shock recovery times, survival, and developmental time. Maternal effects on these traits were often substantial during early offspring development, but these effects either diminished in magnitude or even changed direction as development progressed. In conclusion, our study reveals a dynamic shift in the magnitude and direction of maternal effects on offspring phenotypes in *D. melanogaster*, highlighting the interplay between maternal influence and offspring developmental stage in shaping adaptive responses to environmental variation.

## INTRODUCTION

Phenotypic plasticity allows organisms to respond rapidly, *i*.*e*. within a generation, to environmental changes through adaptations in behavior, physiology, and morphology (Agrawal, 2001; West-eberhard, 1989; Whitman & Agrawal, 2009). For example, in males of the cichlid fish *Astatotilapia burtoni*, the presence of other males triggers a transition from non-aggressive to aggressive behavior within minutes to defend mating partner (Burmeister et al., 2005). Besides this within-generation plasticity, an organism’s phenotype can also be influenced by the parental phenotype and/or environment, a phenomenon called *trans-generational plasticity*. Trans-generationally induced adaptations to potential environmental stressors have evolved many times and are widespread. For instance, parental exposure to mild heat stress can increase the offspring’s ability to cope with heat stress in fish (Donelson, 2015), plants (Whittle,2009), and insects (Lockwood et al., 2017).

The documented impact of both within- and transgenerational plasticity raises the question of how their interplay shapes offspring phenotypes in fluctuating environments where conflictual information between parents and offspring may arise. This question is hard to answer as different sources of environmental cues can affect a given phenotype. For example, in the fruit fly *Drosophila melanogaster* adults respond to an acute heat shock by upregulating the expression of heat shock proteins that increase heat tolerance (Z. W. Li et al., 2009). Heat shock proteins can also be maternally transmitted to the embryo and provide protection against excess heat in the offspring (Lockwood et al., 2017). This shows that heat shock protein-dependent physiological adaptation to heat can be caused both maternally and by the offspring themselves. Such maternal effects can enhance offspring fitness in fluctuating environments. This phenomenon can be regarded as maternal ‘’anticipation’’ of the future environment of their offspring and may increase the chances that the phenotype of their offspring adaptively matches the anticipated environment (e.g., anticipatory maternal effects; Marshall, 2006). However, the extent of maternal effects on offspring phenotype and the factors influencing this impact throughout development remain poorly understood, limiting our understanding of the adaptive nature of these effects. First, adaptive parental effects require that the parental environment is predictive of the offspring environment: in case of a mismatch between the anticipated and the realized environments, parental effects may be disadvantageous. Second, maternal influence on offspring phenotype presumably is adjusted depending on offspring needs during development (Green & McCormick, 2005) and offspring may also become less sensitive to maternal information once they become more capable of perceiving their environment themselves. The more advanced the organisms’ development, the more accurately it may sense its environment and the more unreliable parentally provided information becomes. Organisms may thus be expected to optimize their use of different sources of information regarding their environment by relying on parental information early on and on information acquired by their own sensory system later during development. Indeed, maternal effects on body size are initially high in juveniles of the fish *Poecilia parae* but decline to zero at sexual maturity (Lindholm et al., 2006). However, empirical data that disentangle the relative contribution of parental effects on offspring phenotype at different stages of development are scarce, limiting our understanding of the temporal dynamics of parental effects on offspring phenotype during development.

An excellent characteristic to close this gap is the temperature response in the fruit fly *D. melanogaster*. Temperature is a relevant environmental factor because (i) the metabolism and physiology of ectothermic species, such as flies, strongly vary with temperature, and (ii) temperature fluctuations are omnipresent making it likely that there has been a long history of selection for the ability to induce rapid and plastic adaptions to fluctuating temperatures. Fruit flies have a short developmental time of 10 days at 25 degrees making them ideal to dissect the factors regulating plastic temperature adaptations. Both within-generational (Hercus et al.2003, Hoffman et al.,2003) and transgenerational temperature adaptations (Gibert et al., 2001; Mohan et al., 2018) have been documented in fruit flies. However, the extent to which maternal effects influence offspring phenotype changes during development in this species remains, to our knowledge, unknown. In this study, we used a full factorial design that systematically varied maternal and offspring temperature and measured fitness-related traits, i.e., developmental time, cold shock and heat shock recovery times, and survival at different developmental stages. We quantified the change in magnitude and direction of maternal temperature effects on offspring phenotype as development progressed. We hypothesized that maternal effects on offspring phenotype are larger earlier in development. We further hypothesized that matched environmental conditions between mother and offspring may result in faster recovery, higher survival, and faster development than unmatched conditions between two generations early in development.

## MATERIALS AND METHODS

### *Drosophila melanogaster* strains and rearing

Both the laboratory strain *Oregon-R* and wild-collected *D. melanogaster* flies were used in this study. Wild flies were collected in the province of Groningen, the Netherlands, during the summer of 2017, using banana and dry yeast-baited traps. Male offspring of captured females were used for species identification to eliminate potential *D. simulans* contamination. After identification, 250 *D. melanogaster* females were individually placed in vials to lay eggs. After their eggs hatched, 25 daughters and 25 sons from each female were collected by using CO_2_ anesthesia to mix the F1 generation of those females to create an outbred population. 200 – 250 Flies with an approximately balanced sex ratio were maintained in standard culture bottles filled with 45 ml food (described above). Flies were kept in the lab for 12 generations to adapt to laboratory conditions under 22-23 °C before experimentation.

The reason for using 2 different fly strains was simply due to their availability in the laboratory. The Oregon-R strain used to record cold and heat shock recovery data was a long-term lab-adapted strain. The wild-type flies used for survival and developmental time data were kept in the lab for 12 generations before the experiment. Adult heat and cold shock recovery times were similar in both strains as well as another lab strain *Canton-S* (Kohlmeier et.al, unpublished).

Flies were maintained in 177 ml, polypropylene culture bottles filled with 45 ml food (agar (10 g/l), glucose (167mM), sucrose (44mM), yeast (35 g/l), cornmeal (15 g/l), wheatgerm (10 g/l), soya flour (10 g/l), molasses (30 g/l), propionic acid (5ml of 1M) and tegosept (2 g in 10ml of ethanol)) in the lab temperature (22-23 °C). After two weeks, adults were transferred to new bottles with fresh food for egg laying and removed after 24 hours to avoid larval overcrowding. Twenty-four hours after removing the adults, freshly hatched larvae were collected using a sieve, tap water, and a brush and used for split brood experiments at laboratory temperature.

### Split brood design

To disentangle the effects of parental and offspring temperature on offspring phenotypes, a 2×2 fully factorial design was two different temperatures, 28 and 18 °C were used as maternal and offspring ambient temperatures to maintain two matched and two unmatched groups, designated as HH, CC, and HC, CH, respectively (H for “hot”; 28 °C and C for “cold”; 18 °C, the first letter represents the maternal temperature, and the second letter represents the offspring temperature as in Figure 1). Females were split into two groups after eclosion and individually kept in vials (2.3×9.3 cm) with fly food at hot or cold temperatures. After 24 hours, two freshly eclosed males from the same population kept at room temperature, 22-23 degrees (to mitigate paternal effects in the results) were added to each individual female vial for mating. After 24 hours, H-females were placed individually on a 35 × 10 mm egg-laying patch, fly food supplemented with a dab of yeast paste (1g yeast/ml water), covered by an empty vial on top. The same step was done with the females in C after 48 hours instead of 24 hr due to the delaying effects of cold temperatures on maturity. Females were transferred to a new food patch every 2h. The eggs laid on their former patches were randomly assigned to either cold or hot treatment and transferred to a fresh food patch. The split brood design and following assays were conducted simultaneously in two different chambers, one maintained at 28°C and the other at 18°C. Both chambers operated on a 12:12 day-night light cycle from 9 am to 9 pm.

**Figure 1:**
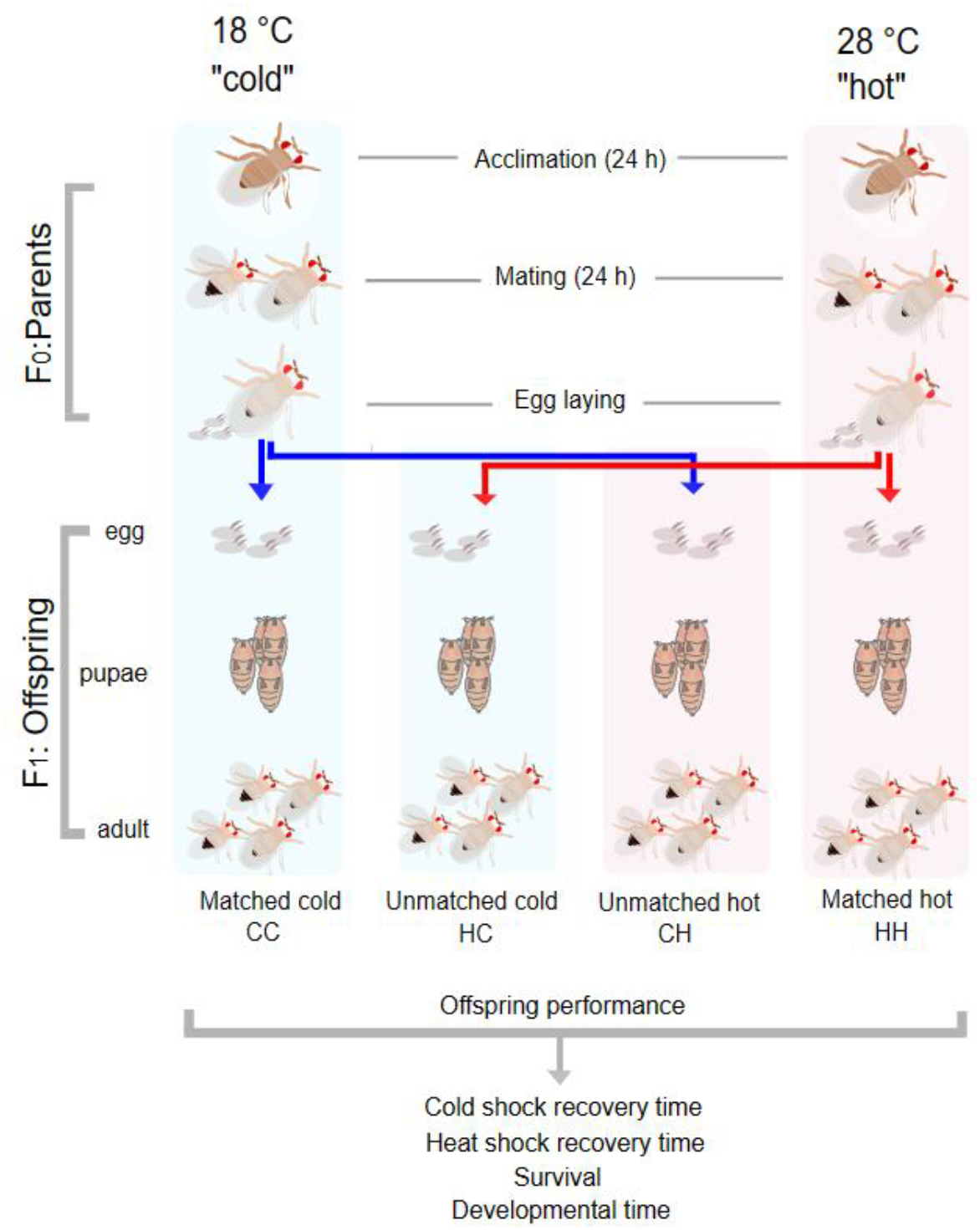
Schematic of the 2 × 2 fully factorial split brood design for estimating parental and offspring temperature effects on offspring performance. On the 1^st^ day after eclosion, each female from the parental generation experienced either 18 or 28 °C for 24 hours. On the 2^nd^ day, they mated with males that had experienced the same conditions for 24 h at 28 °C and 48 h at 18 °C. Afterward, females were separated from males to lay their eggs. Every 2 h, the clutches laid by individual females were randomly split in two and each half was transferred to either 18 or 28 °C for further development. The resulting four groups of offspring experienced temperatures that either matched or mismatched the temperatures of their parents and were subsequently tested for their performance during early development or in adulthood. Cold and heat shock recovery times (from 175 and 144 unique mother IDs) were measured in L1 larvae (n=380, n=246) and adults (n=124, n=127) Survival and developmental time (from 1943 and 1956 unique mother IDs) were measured from the egg stage until the pupal stage(n=7695) and until adulthood (n=7695).

### Heat shock recovery time assay

To quantify larval heat shock recovery times, L1 larvae were placed in small glass vials (40 mm × 8 mm × 0.9 mm) filled with 0.7 ml black agar gel (1% agar, 1% activated charcoal) to make the larvae visible. These vials were placed in larger vials which were attached to a Styrofoam floater. The vials were then floated in a water bath heated to 40.5°C for 30 minutes. After heat exposure, the larvae were monitored under a stereomicroscope until the first bodily movement, which usually occurred in the rostral part of the larvae. Recovery time was defined as seconds passed between the end of the heat exposure and the first bodily movement observed. To quantify heat shock recovery in adult flies, 6 freshly enclosed flies (3 females, 3 males) per mother were aged in a common garden at room temperature (23 °C) for 5-6 days to prevent direct temperature effects in adulthood. Flies were transferred individually into small glass vials (40 mm × 8 mm × 0.9 mm), closed with cotton plugs, and submerged in a 40 °C water bath for 7 minutes and 20 seconds. Flies were then individually transferred to empty Petri dishes (35 × 10 mm) by a paintbrush at 23 °C and recovery time, defined as the number of seconds until flies stood on their legs, was recorded. The reason for different heat exposure duration and temperatures during larval and the adult stages is that no larvae were knocked out after exposure to the adult heat shock protocol. Thus, we ensured that all individuals (both larvae and adults) that were exposed to heat shock were paralyzed but not killed.

### Cold shock recovery time assay

Each L1 larva was placed on a 94-well PCR plate with the wells pre-filled with 0.7 ml black agar gel (1% agar, 1% activated charcoal) to prevent the larvae from sticking to the bottom of the wells and to make the larvae visible. Afterward, the plate was embedded in ice for 4 hours. To quantify cold shock recovery in adults, flies were kept at 0 °C for 5 hours and 20 minutes in an ice-filled box. After cold exposure in adults, the same protocol for adult heat shock recovery was used for measuring recovery time.

Both cold and heat shock recovery time assay protocols were adjusted from adult flies to larvae to make them effective for larvae. Throughout these modifications, we ensured that the average duration from heat or cold exposure until the flies were immobilized remained consistent between larvae and adults.

### Developmental time and survival assay

Developmental time was defined as the number of hours passed from clutch splitting (a few hours after eggs were laid) until pupation or emergence. The number of freshly emerged pupae, adults, and their developmental time were quantified by counting them three times per day, at 9 am, 2 pm, and 6 pm (as three observation times), until eclosions were completed. The emergence of fruit flies from pupae primarily occurs during the daytime, with a notable peak around dawn (De et al., 2012). Therefore, we constrain our observations to the period between 6 pm and 9 am.

The developmental time of the pupae or emerging adults was recorded as the latest observation time. Each observation lasted around 45 min. This leads to a couple of hours of error in estimating the exact emerging time. Therefore, an error of up to 4,5 hours occurred during the day (between 9 am and 1:30 pm or 1:30 pm and 6 pm).

Survival was calculated on a per-mother basis as the proportion of eggs reaching the pupal or adult stage.

### Statistical Analysis

All analyses were conducted with R (version 4.3.3; R Core Team 2023) in the RStudio development environment (version Ocean Storm; RStudio Team 2022). To fit fully Bayesian models, we used the brms package (version 2.14.0; Bürkner, 2017, 2018), interfaced with the Stan MCMC sampler via the cmdstanr package (rstan version 2.32.6; Stan Development Team 2022). All figures were produced with the ggplot2 package (Wickham, 2016).

#### Models

For the three non-negative “duration” variables heat shock recovery time, cold shock recovery time and developmental time we fitted lognormal models of the form:

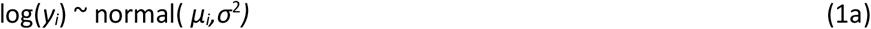

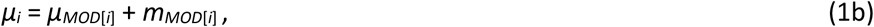

where the index *i* refers to an individual. *MOD*[i] denotes an individual’s state out of the 8 possible combinations of maternal temperature (*M* ∈ {C,H}), offspring temperature (*O* ∈ {C,H}) and developmental stage (*D* ∈ {Early,Late}). Thus, the *μ*_*MOD*[*i*]_ comprise the “fixed” part of the model, the means for each of the 8 combinations of the three binary predictor variables. This model is equivalent to a model with main effects of *M, O* and *D*, all three two-way interactions *M* × *O, M* × *D, O* × *D*, and the three-way interaction *M* × *O* × *D*. Indeed, this is how we specified the model in the R code, using treatment contrasts.

The term *m*_*MOD*[*i*]_ refers to a “random” effect for the mother of individual *i*. Since we used a split brood design, where for each mother the offspring were equally split between treatments *O* = C and *O* = H, we allowed these maternal effects to depend on *O* and we estimated maternal effect variances and covariances according to

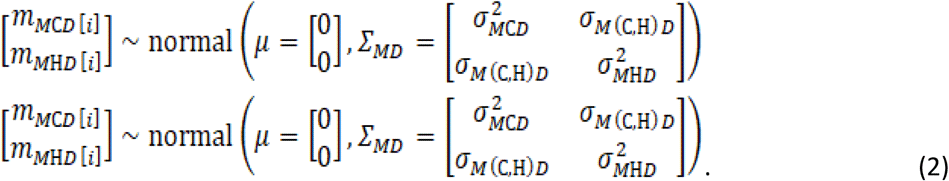

Hence, we estimated four covariance matrices *∑_MD_*, for each of the four combinations of maternal treatment *M* and developmental stage *D*.

For the survival data we fitted logistic models of the form

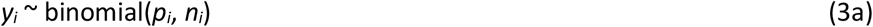

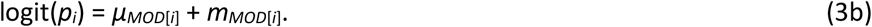

Here the analyses are at the maternal level. Thus, *y*_*i*_ is the number of surviving offspring of mother *i*, out of an initial number of *n*_*i*_ offspring, and are modeled as draws from a binomial distribution with parameter *p*_*i*_ that specifies the probability of survival. The logit transform log(*p*_*i*_/(1−*p*_*i*_)) constructs the linear predictor. The interpretations of the fixed and random effects are similar to the above lognormal models.

#### Effect sizes

For the lognormal models we report dimensionless effect sizes as ratios on the original scale (median and 95% highest posterior density [HPD] credible intervals [CIs]), while for the logistic models we report odds ratios. We used the emmeans package (Lenth 2023) to calculate these effect sizes and corresponding CIs. For the logistic models we used the emmeans package’s bias correction methods (delta method) to correct for biases due to the non-linearity of the logit transform.

Specifically, effects of maternal temperature *M*, for a given offspring temperature *O* ∈ {C,H} and developmental stage *D* ∈ {E,L} are quantified as C/H ratios:

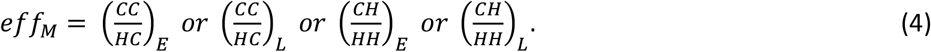

For example, the first ratio on the right of the equation is the effect of cold relative to hot mothers for cold offspring during the early developmental stage (with L meaning late, i.e. the adult stage).

Likewise, the effects of offspring temperature are reported as

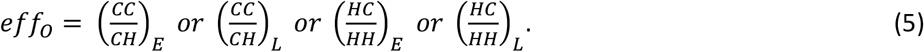

Relative effects of maternal temperature for C offspring and H offspring, given a developmental state *D* are given by ratios of ratios:

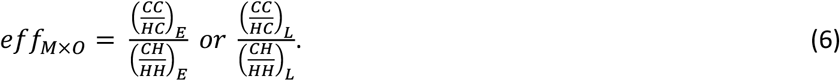

These correspond to the two-way interactions *M* × *O* within developmental stages.

Relative effects of maternal temperature for E(arly) and L(ate) developmental stages, given an offspring temperature *O*, are reported as

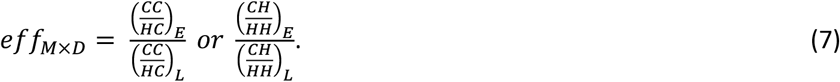

These correspond to the two-way interactions *M* × *D* within offspring temperatures.

We do not report on the remaining two-way interaction *O* × *D*, as this is of little interest to us.

The three-way interaction is reported as a ratio of a ratio of a ratio:

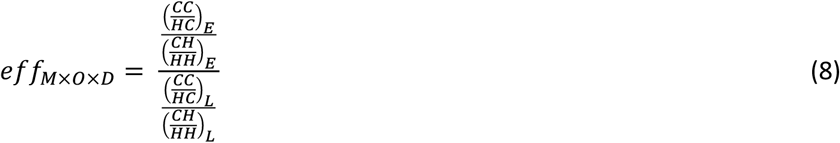

#### Operational definitions of maternal effects

We can distinguish between several kinds of maternal effects in our analysis. First, the average effects of maternal temperature *M* for a given offspring temperature *O* and developmental stage *D*. We refer to these as the *conditional maternal effects*. These are the “fixed” effects in the models (1ab) and (3ab), and they are quantified as C/H ratios according to (4). Within-developmental stage comparisons between conditional maternal effects for C and H offspring are quantified by the ratio (6). Especially relevant for this study are the within-offspring temperature and between-developmental stage comparisons, as quantified by the ratios (7).

Second, there are the “random” maternal effects, given by the *m*_*MOD*[*i*]_ terms in models (1ab) and (3ab). These we call the *individual maternal effects*. For the purpose of this study, these are of less interest than the conditional maternal effects. Their importance is quantified by the variances 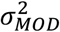 in (2).

Third, there are the correlations in trait values between C and H offspring from the same mothers, as quantified by the correlation coefficients.

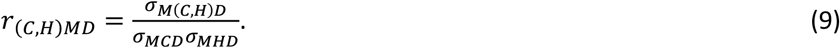

We call these the *correlational maternal effects*.

#### Hypothesis testing

To test hypotheses regarding effect sizes, we used the probability of direction, pd (Makowski et al., 2019), which is the posterior probability of the sign of the posterior median. More specifically, the pd-value equals the proportion of the posterior density that has the same sign as the median of the posterior density; this value can be thought of as a rough Bayesian equivalent of the frequentist p-value. For example, if the estimated posterior median of an effect size is positive, the corresponding pd value is the posterior probability that the effect size is positive. We use stars (*) to indicate the “significance” level of pd values: * = pd > 0.9091 ((odds) ratios 10:1), ** = pd > 0.9901 ((odds) ratios 100:1), *** = pd > 0.9990 ((odds) ratios 1000:1). For the ratios (4)-(8) the pd values refer to posterior probabilities that the ratios are smaller or larger than 1.

#### Priors

Regarding priors, for “fixed” effects we used “weakly informative” (Lemoine, 2019) priors. Specifically, for intercepts we used normal (0,5) and for contrasts normal (0,1). For group-level (“random”) effects we used the default priors of brms, i.e., a half t-density with 3 degrees of freedom for standard deviations and a LKJ (1) density for correlations.

#### MCMC details and diagnostics

For each model we ran 4 chains with 1,000 warmup iterations, followed by 2,500 sampling iterations thus creating 10000 posterior samples per model. Proper mixing of chains was monitored with trace plots and convergence of chains by verifying that Rhat values were close to 1. Model fits were also evaluated by inspecting posterior predictive checks, using the pp_check () function of brms.

### Additional tools and software

Our textual data were analyzed using a natural language processing tool, ChatGPT (OpenAI, 2024). Graphical representations were created using vector graphics software, Inkscape (Inkscape, 2022).

## RESULTS

### Cold shock recovery time

Offspring temperature influenced cold shock recovery time (CSRT), both at the larval stage and the adult stage (Figure 2A and Table 1). In addition, flies from H-mothers reared under hot (H) conditions had significantly longer recovery times than flies reared under cold (C) conditions, but this was not the case for larvae from C-mothers. Indeed, the interaction between maternal temperature and offspring temperature was significant (OxM|D=E, pd=0.99). There was also a significant 3-way interaction OxMxD (pd=0.97), indicating that the OxM interaction declined from the larval to the adult stage (Figure 2A and Table 2).

**Table 1:**
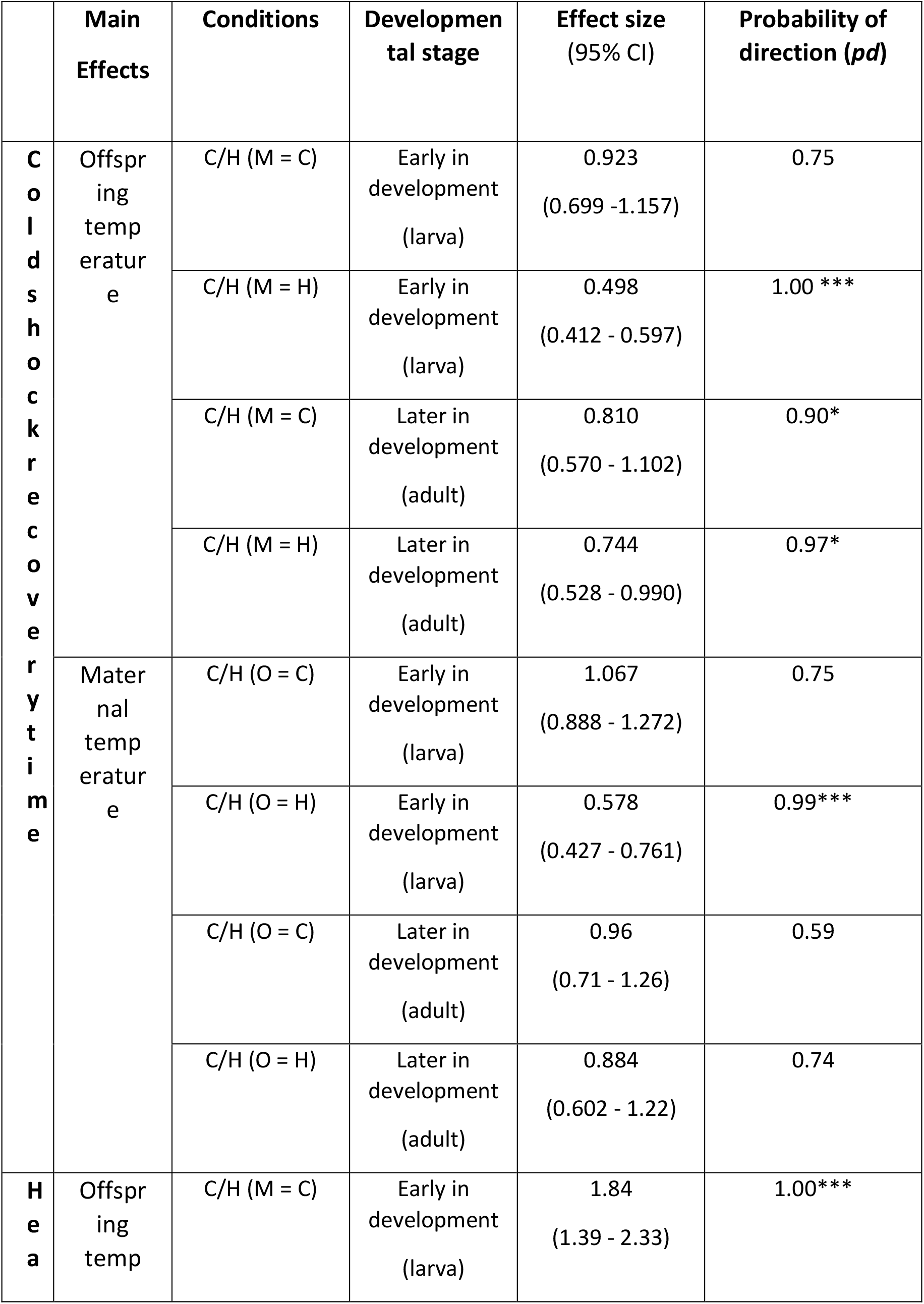
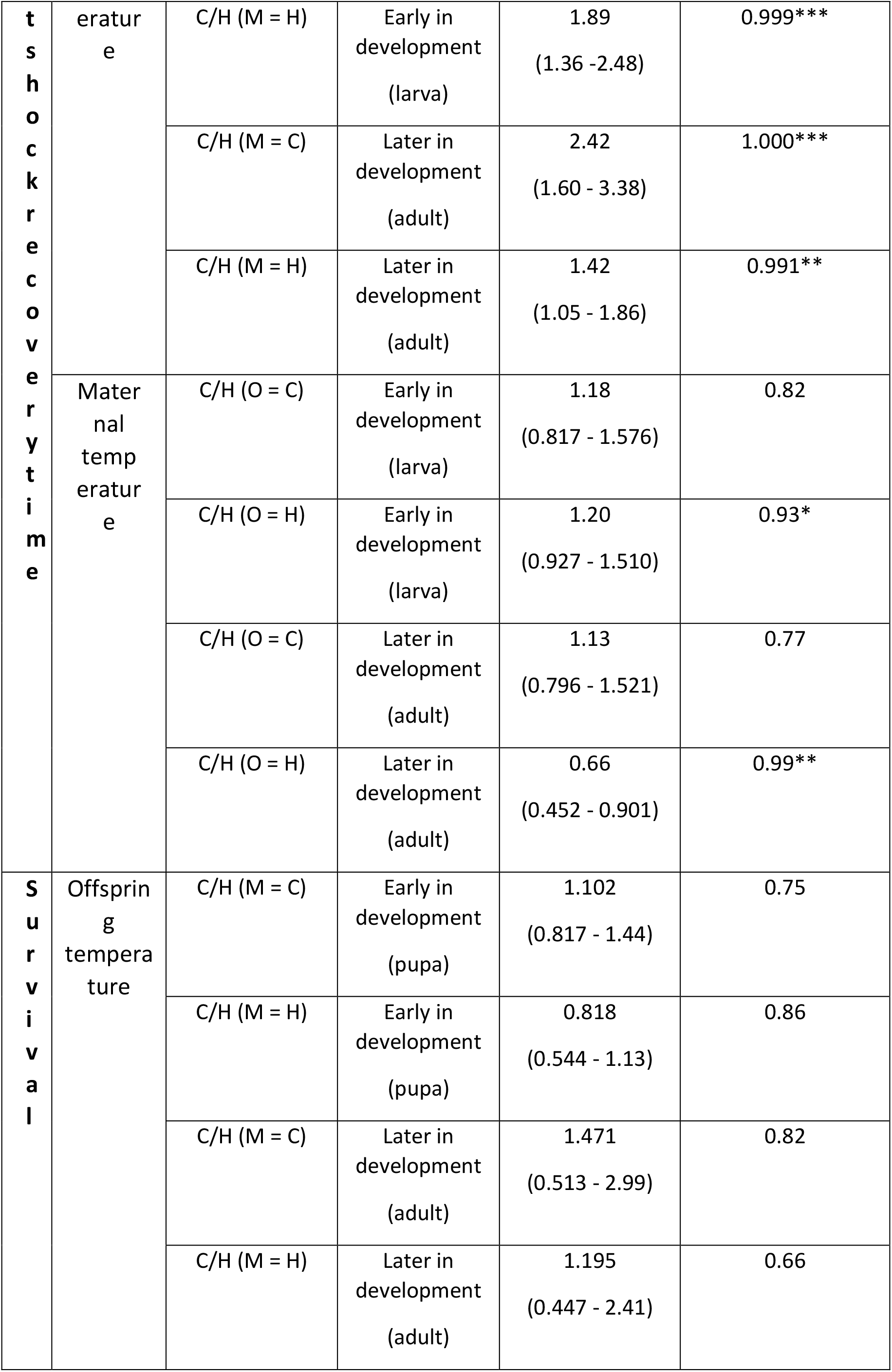
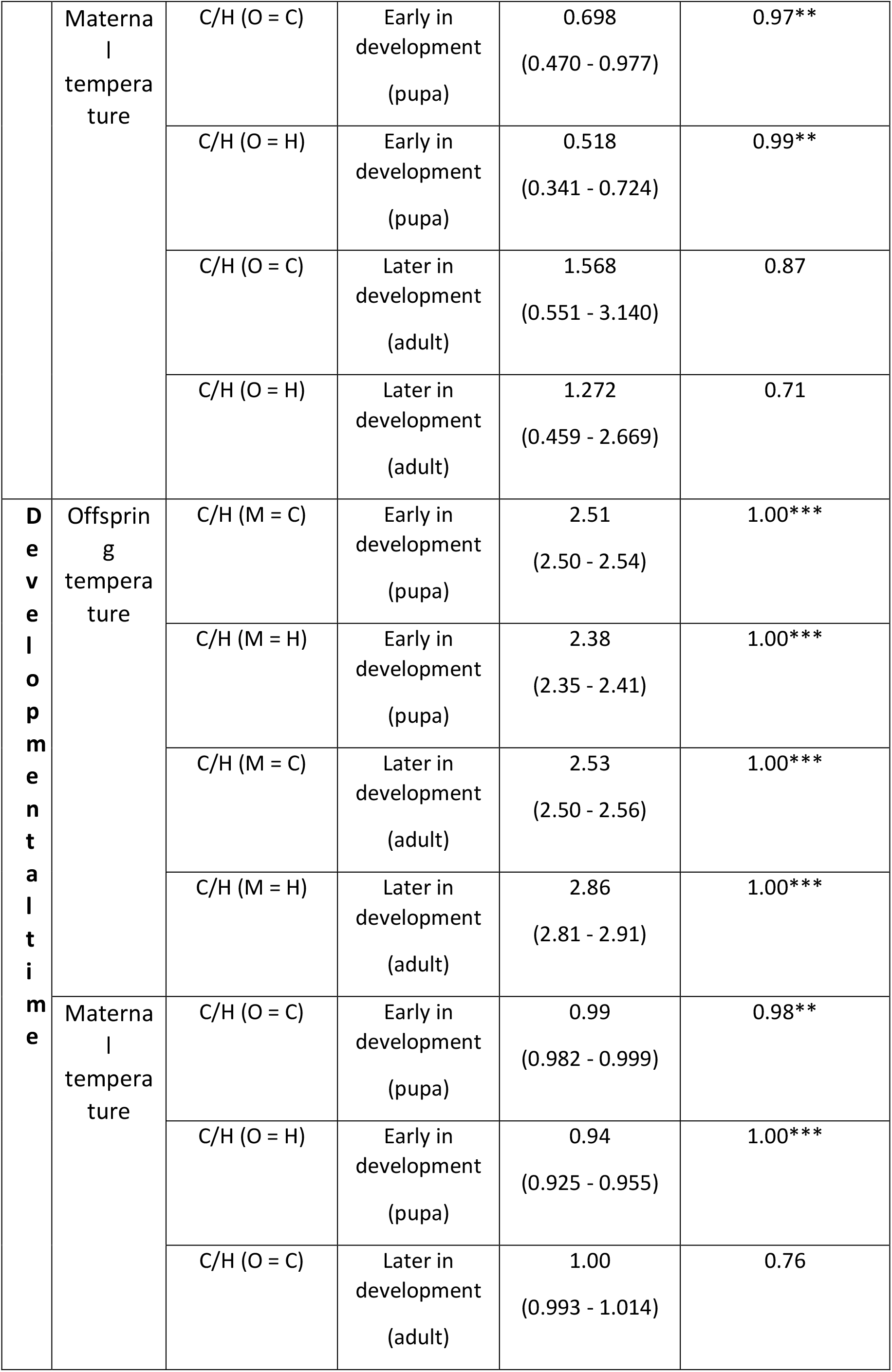
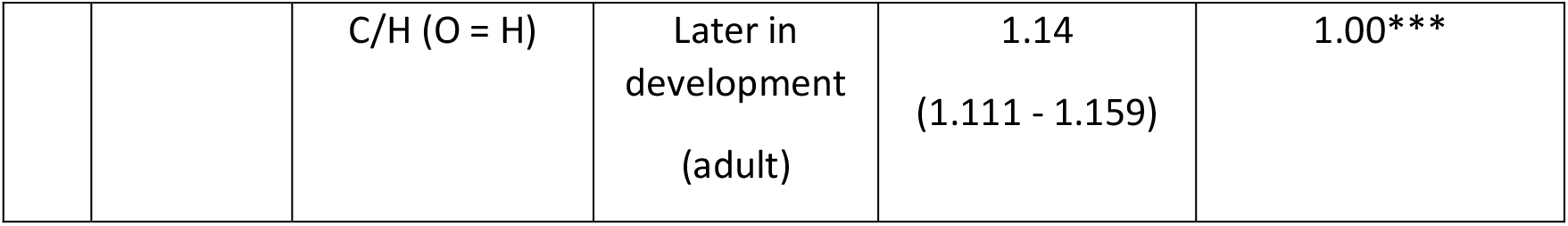
Overview of the main effects of the Bayesian analysis. The “Main effects” column represents the predictor variables used, offspring temperature (O), and maternal temperature (M). The “Conditions” column represents the subgroups that arose due to the split brood design. “C” defines cold, and “H” defines hot conditions. For instance, “M=C” was used to define the condition when the maternal temperature is cold. The ”Developmental stage” column defines early and later development groups, respectively larva or pupa and adult. The “Effect size” column provides a measure of the magnitude/strength of the effects observed from each main effect. The “pd” column provides pd values associated with each effect. (* pd > 0.909 or 10:1 odds, ** pd > 0.990 or 100:1 odds, *** pd > 0.999 or 1000:1 odds).

**Table 2:** Overview of interaction terms: 4 different types of 2-way interactions and 3-way interaction. The first one is for describing the developmental stage effect on maternal effects in both C and H condition. The second type of 2-way interaction is for describing offspring temperature on maternal effects (M*O) in early (E) and later (L) in development. 3-way interaction to describe both offspring temperature and developmental stage effects on maternal effects (M*O*D). The “*pd*” column provides values associated with each effect. Pd values are used to assess the significance of the effects we have found (* = pd > 0.9091, ** = pd > 0.9901, *** = pd > 0.9990).

| Effects/Traits |  | Cold shock recovery time | Heat shock recovery time | Survival | Developmental time |
| --- | --- | --- | --- | --- | --- |
| Interactions | Effects | Probability of direction (*pd*)<br>Effect size (95% CI) |  |  |  |
| M*D \| O=C | (CC/HC)_E / (CC/HC)_L | **0.72**<br>1.10<br>(0.75 - 1.49) | **0.56**<br>1.04<br>(0.62 - 1.56) | **0.97 \***<br>0.44<br>(0.15 - 0.90) | **0.97\***<br>0.98<br>(0.97 - 1.00) |
| M*D \| O=H | (CH/HH)_E/ (CH/HH) | **0.97\***<br>0.64<br>(0.39 - 0.97) | **0.99 \*\***<br>1.83<br>(1.12 - 2.62) | **0.97 \***<br>0.40<br>(0.12 - 0.87) | **1.00\*\*\***<br>0.82<br>(0.80 - 0.84) |
| M*O \| D=E | (CC/CH)/(HC/HH) | **0.99 \*\*\***<br>1.85<br>(1.31 - 2.45) | **0.55**<br>0.97<br>(0.62 - 1.38) | **0.90**<br>1.35<br>(0.83 - 2.01) | **1.00\*\*\***<br>1.05<br>(1.03 - 1.06) |
| M*O \| D=L | (CC/CH)/(HC/HH) | **0.64**<br>1.09<br>(0.65 - 1.62) | **0.98 \***<br>1.70<br>(1.05 - 2.63) | 0.64<br>1.21<br>(0.29 - 2.90) | **1.00\*\*\***<br>0.88<br>(0.86 - 0.90) |
| M*O*D | (CC/CH)/(HC/HH)_E<br>/(CC/CH)/(HC/HH)_L | **0.97 \***<br>1.69<br>(0.93 - 2.75) | **0.97 \***<br>0.57<br>(0.28 - 0.94) | **0.57**<br>1.10<br>(0.30 - 2.82) | **1.00\*\*\***<br>1.19<br>(1.16 - 1.22) |

**Figure 2:**
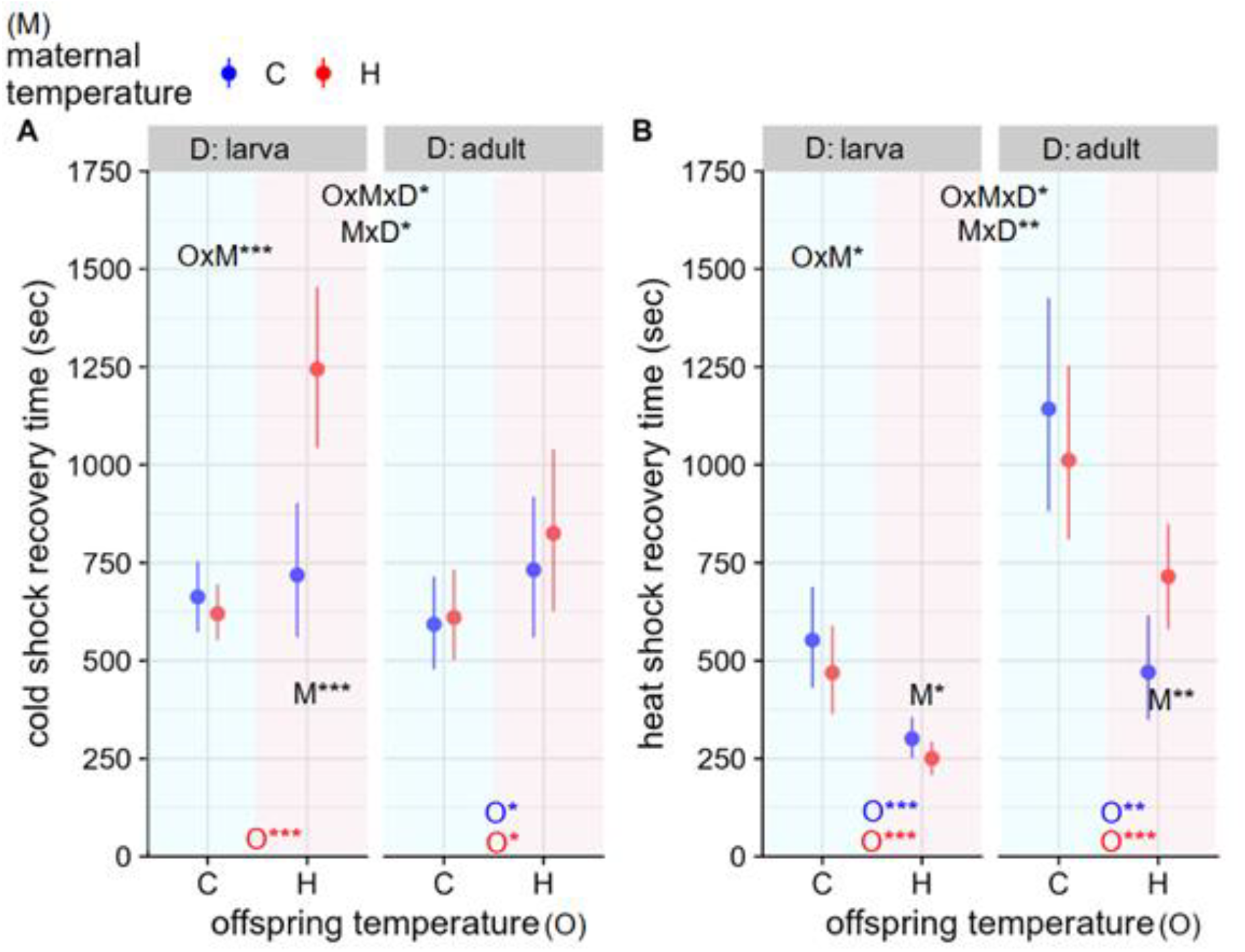
The effects of maternal and offspring temperatures on cold shock and heat shock recovery times in larval and adult stages. C and blue denote 18°C while H and red denote 28°C. Dots indicate posterior medians and error bars 95% credible intervals. Stars indicate “significant” differences between groups according to the probability of direction (* pd>0.909 or 10:1 odds; ** pd>0.990 or 100:1 odds; *** pd>0.999 or 1000:1 odds). M refers to maternal effects, i.e. within offspring temperature ratio of C_mother_/H_mother_ or blue/red. O refers to offspring effects, i.e. within the maternal temperature ratio of C_offspring_/H_offspring_. The color O indicates the maternal temperature, i.e. blue O refers to the offspring from cold mothers. MxD refers to maternal effects*developmental stage interactions. MxO refers to the within-developmental stage maternal temperature*offspring temperature interaction. OxMxD refers to the 3-way maternal temperature*offspring temperature*developmental stage.

The only significant effect of maternal temperature occurred in larvae raised in H-conditions, where larvae of H-mothers had a considerably longer CSRT than larvae of C-mothers (pd=0.99, Figure 2A, Table 1).

Between the larval stage and the adult stage, the effect of maternal temperature differed for H-offspring (MxD|O=H, pd=0.97), indicating a decline in the magnitude of a conditional maternal effect from the larval to the adult stage.

### Heat shock recovery time

Offspring temperature had an effect on heat shock recovery time (HSRT) both at the larval stage and the adult stage (Figure 2B and Table 1). Flies reared under H conditions always recovered faster from heat shock than flies reared under C conditions, regardless of maternal temperature and developmental stage. However, there were some conditional maternal effects as well: H-larvae of H-mothers recovered even faster than H-larvae of C-mothers (pd=0.93, Figure 2B and Table 1). For adults, this maternal effect had the opposite direction: H-adults of C-mothers recovered faster than H-adults of H-mothers (pd=0.99, Figure 2B and Table 1). Indeed, for H-flies, the effect of maternal temperature differed significantly between larval and adult stages (MxD|O=H, pd=0.99), indicating a change of direction in a maternal effect from the larval to the adult stage.

Additionally, during the adult stage, the effect of maternal temperature differed between C-flies and H-flies (OxM|D=L, pd=0.98), such that the effect of maternal condition was larger and opposite in flies reared in hot conditions.

There was also a significant 3-way interaction OxMxD (pd=0.97), indicating that the MxD interaction varied between C-flies and H-flies (Figure 2B and Table 2).

### Survival

Temperature experienced during development did not significantly affect offspring survival, both from egg until pupation and from pupal to adult stage (Figure 3A and Table 1). However, the maternal temperature did affect survival until the pupal stage: larvae of H-mothers survived better than larvae of C-mothers (pd=0.99, Figure 3A and Table 1), irrespective of offspring conditions. During the pupal stage, survival seemed unaffected by maternal temperature. This difference between stages in the effect of maternal temperature was itself significant (MxD|O=C and MxD|O=H both pd=0.97). Thus, regardless of offspring temperature, maternal effects on survival declined from the pupal to the adult stage (Figure 3A and Table 2). No other significant interactions were observed.

**Figure 3:**
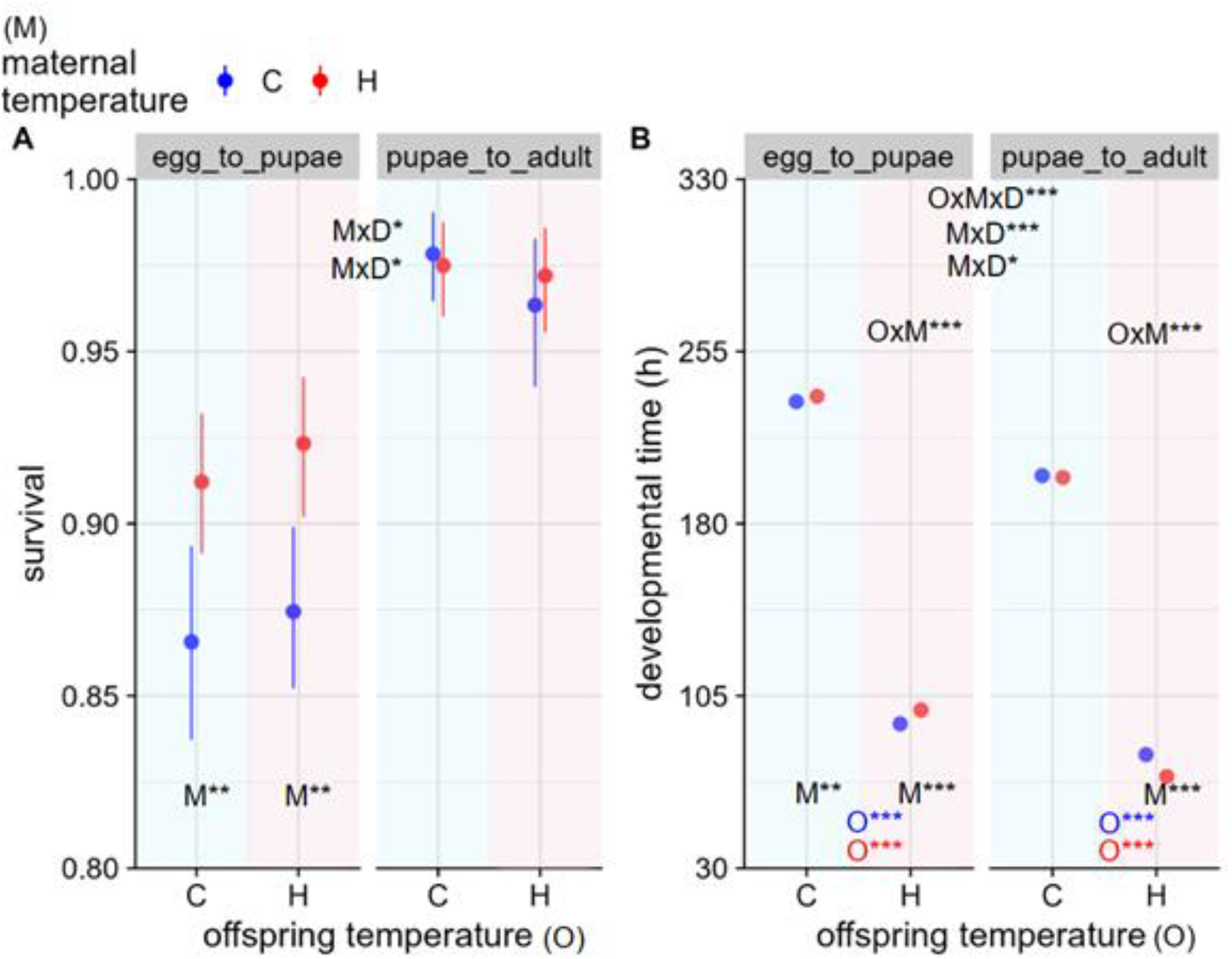
The effect of maternal and offspring temperature on survival and developmental time in both early and late development; larval and adult stages. C and blue denote 18°C while H and red denote 28°C. Dots indicate posterior medians and error bars 95% credible intervals (they are too narrow relative to the variability in developmental time data). Stars indicate “significant” differences between groups according to the probability of direction (* pd>0.909 or 10:1 odds; ** pd>0.990 or 100:1 odds; *** pd>0.999 or 1000:1 odds). M refers to maternal effects, i.e. within offspring temperature ratio of C_mother_/H_mother_ or blue/red. O refers to offspring effects, i.e. within the maternal temperature ratio of C_offspring_/H_offspring_. The color O indicates the maternal temperature, i.e. blue O refers to the offspring from cold mothers. MxD refers to maternal effects*developmental stage interactions. MxO refers to the within-developmental stage maternal temperature*offspring temperature interaction. OxMxD refers to the 3-way maternal temperature*offspring temperature*developmental stage.

### Developmental time

Offspring ambient temperature had a strong effect on developmental time (DT) both from egg until pupation and from pupal to adult stage (Figure 3B and Table 1). Flies reared under H conditions developed more than twice as fast as flies reared under C conditions.

Maternal effects on DT occurred during both the larval and pupal stages. When taken together, larvae raised under C and H conditions developed faster when they had C-mothers. (pd=1.00, pd=1.00 Figure 3B and Table 1). For C-pupae, no significant maternal effect was observed while in H-pupae maternal effects had changed signs from the larval stage: pupae of H-mothers developed faster than pupae of C-mothers (pd=1.00, Figure 3B and Table 2). For both C-pupae and H-pupae, the maternal effects had changed direction significantly compared to the larval stage (MxD|O=C, pd=0.97; MxD|O=H, pd=1.00).

All other interactions were also significant. During both developmental stages, the maternal effects were of greater magnitude for H-flies than for C-flies (OxM|D=E and OxM|D=L, both pd=1.00). The 3-way interaction OxMxD (pd=1.00) indicates that these 2-way interactions changed direction between developmental stages (Figure 3B and Table 2).

## DISCUSSION

We found that maternal temperature effects on offspring phenotype often changed substantially during development in *D. melanogaster*. Maternal effects influenced offspring performance with varying strength between early and later in development, including heat shock and cold shock recovery times, as well as survival and developmental speed. These maternal effects tended to be greater at early than later developmental stages and sometimes even reversed in sign in adulthood. Among the 8 conditions (4 traits by 2 offspring temperatures) tested, 4 showed a decrease in maternal effects magnitude, while only 2 showed a significant increase yet reversed in direction (Table 3). Similarly, a reduction in maternal effects was found in different fish, *Acanthochromis polyacanthus* (Donelson et al., 2008) and *Poecilia parae* (Lindholm et al., 2006).

**Table 3:** The summary of magnitude and direction change in maternal temperature effects from early to late development. The words “decreased” and “increased” were used to explain the change in the magnitude of maternal effects, while “reversed” was used to explain to change in the direction of maternal effects from early to late in development. Otherwise “-” was used to indicate no significant change both in magnitude and direction.

| Offspring Trait | C-offspring |  | H-offspring |  |
| --- | --- | --- | --- | --- |
|  | Maternal effects change |  |  |  |
|  | in magnitude | in direction | in magnitude | in direction |
| Cold shock recovery time | - | - | decreased | - |
| Heat shock recovery time | - | - | increased | reversed |
| Survival | decreased | - | decreased | - |
| Developmental time | decreased | - | increased | reversed |

In addition to varying in magnitude, some maternal temperature effects also varied in sign over developmental time. This might be best explained in the light of the different benefits of maternal effects on mothers and their offspring (Marshall & Uller, 2007). Indeed, anticipatory maternal effects boost both maternal and offspring fitness; selfish effects prioritize maternal fitness over offspring fitness; bet-hedging effects minimize variance in maternal fitness by producing offspring with varied phenotypes; lastly transmissive effects decrease both maternal and offspring fitness. Anticipatory maternal effects occur when the environmental conditions for both parents and offspring match, allowing mothers to “anticipate” the future conditions for their offspring. This maximizes maternal reproductive success by enhancing offspring fitness. Although evidence for such effects is scarce, they have been detected in plants (Galloway & Etterson, 2007; Herman & Sultan, 2011), vertebrates (Salinas et al., 2013), and invertebrates (Cahenzli & Erhardt, 2013). Traits measuring offspring performance under stressful temperatures, offspring cold shock and heat shock recovery time, showed an interesting pattern in early development, by some means explained by the core logic of anticipatory maternal effects: for the cold shock recovery time of H-larvae (developed in the **H**ot condition), offspring of C-mothers (experienced the **C**old condition), despite of the unmatched conditions recovered faster than offspring of H-mothers (matched). Perhaps offspring conditions (H) do not match maternal conditions (C), but the cold shock treatment itself “does’’ match maternal conditions. For the heat shock recovery time, H-larvae recovered faster had H-mothers, presumably advantageous for coping with extreme heat. Importantly, this direction of maternal temperature effects as a boost to faster recovery in early development, regardless of matching status, disappeared in adulthood.

In some cases, one particular maternal environment always leads to “the best” offspring performance regardless of the offspring’s environment. Such effects are called *silver spoon effects* and benefit offspring regardless of environment matching between offspring and mothers (Engqvist & Reinhold, 2016). During the larval stage, the offspring of H-mothers had higher survival than the offspring of C-mothers, regardless of the offspring’s thermal regime. However, this effect was not observed at later developmental stages, as neither maternal nor offspring temperature affected survival in the adult stage. This outcome suggests both a decline in such maternal effects and strength in survival as development progresses. However, adult mortality was generally so low that detecting maternal effects during this stage might have been limited by the low sample size. As another silver spoon effect, larvae of C-mothers developed faster than larvae of H-mothers, regardless of their temperature, and this pattern reversed during the pupal stage. As a result, silver spoon effects were “lost” due to the change in the direction of maternal effects from early development to adulthood in developmental time. The combination of these outcomes, along with the varying magnitude and direction of maternal effects on specific traits during development, highlights the complex nature of maternal temperature effects. Understanding the factors that cause the varying influence of maternal effects on offspring fitness and performance during ontogeny can reveal potential evolutionary implications of these effects.

Maternal effects only manifest themselves in the offspring phenotype if the offspring are sensitive to maternal information. Offspring might become less sensitive to maternal environmental information as they can act on the information they gather themselves. This may result in weaker maternal effects. Our results suggest that maternal information may become less relevant as offspring begin to sense their environment independently. This allows offspring to disregard maternal information (Kuijper & Hoyle, 2015). Moreover, offspring continuously gather environmental information throughout their lives, leading to constant updates to their phenotype (Groothuis & Taborsky, 2015). However, the processes by which offspring select the weight of maternal and its own environmental information to shape their phenotype remains unknown. It is evident that not all environmental information is relevant, and its significance may vary during different developmental stages (Lhomme et al., 2020). Theoretical models suggest that selection favors multiple periods of heightened plasticity throughout an organism’s lifetime (reviewed by Fawcett & Frankenhuis, 2015), indicating that organisms may collect environmental information at various stages before integrating it into their phenotype (Groothuis & Taborsky, 2015). This implies that in our study both offspring and maternal temperature information could contribute to an updated phenotype if maternal information is reliable. Offspring strategy to use maternal information to shape its phenotype includes considering the costs of parent information to the offspring, as well as constraints on information acquisition or the evolution of counter-responses to paternal strategies (Uller & Pen, 2011), on top of the differences in these factors during development. Empirical support for a change in the direction of maternal effects throughout offspring development is limited. As our results demonstrate both a decrease in the magnitude and a shift in the direction of maternal temperature effects over offspring age, we think that such effects can be the result of cost and constraints during maternal information transfer and /or usage.

Maternal effects during development tend to occur under both favorable and stressful conditions (Rösvik et al., 2020), which may facilitate adaptation to novel environments (Fox & Savalli, 1998; Leftwich et al., 2019) or buffer against changing conditions through transgenerational phenotypic plasticity (Shama et al., 2014). We found that maternal effects, besides being influenced by the offspring’s developmental stage, can also depend on other factors such as the offspring environmental conditions. Our study revealed that maternal effects in 75% of the measured traits were dependent on offspring temperature, especially during early development. Specifically, a warmer offspring condition appears to be notably more conducive to the manifestation of maternal effects compared to a colder offspring environment, as observed in both cold shock and heat shock recovery times. Furthermore, the duration of maternal effects can also be dependent on the offspring’s condition. In line with our findings, maternal nutrition effects varied across different environments in soil mites, maternal effects in egg size predominantly influenced increasing variation across generations in high-food environments, while they were weak and decreased over time under low-food environments (Plaistow et al., 2006). Moreover, maternal effects can manifest differently across various traits. Our study revealed differences in both the magnitude and direction of maternal effects among performance and fitness traits. Therefore, interpreting context-dependent maternal effects necessitates conducting multi-trait, multi-environment studies. Univariate life history measures in single environments may underestimate the full significance of intergenerational effects (Plaistow et al., 2006).

This study offers compelling evidence that maternal ambient temperature in *D. melanogaster* can profoundly affect offspring fitness and performance throughout development, influencing factors such as recovery time from heat and cold shock, survival, and developmental time. Furthermore, our research demonstrates that the strength and direction of maternal temperature effects typically shift as offspring progress from early development to adulthood. The overall trend of diminishing maternal effects as development progresses likely stems from offspring becoming less sensitive to maternal information and relying more on current environmental information to determine the phenotype under environmental heterogeneity.

## AUTHOR CONTRIBUTIONS

Pinar Kohlmeier, Jean-Christophe Billeter, Ido Pen and Ton G.G. Groothuis conceived the framework and designed the methodology; Pinar Kohlmeier and Bart Van Schaik (partly) collected the data; Pinar Kohlmeier and Ido Pen analyzed the data; Pinar Kohlmeier, Ido Pen, Ton G.G. Groothuis and Jean-Christophe Billeter led the interpretation and Pinar Kohlmeier wrote the manuscript with the help of JCB, IP and TG. All authors contributed critically to the drafts and gave final approval for publication.

## ACKNOWLEDGEMENTS

We thank Mario Santos Mira for contributing to data collection by sharing adult recovery time data. We thank Philip Kohlmeier for his suggestions to make the manuscript better. Lastly, we thank Gerard Overkamp for his help that made parts of the experiments run efficiently.

## FUNDING INFORMATION

Pinar Kohlmeier was supported by Adaptive Life grants awarded by the University of Groningen to IP, TGGG, and JB as well as the Netherlands Organization for Scientific Research (NWO) ALWOP.279 to IP.

## CONFLICT OF INTEREST

No competing interest to be declared.

## Notes

### Competing Interest Statement

The authors have declared no competing interest.

