## Supplemental Tables for "Temperature-modulated maternal effects vary with offspring developmental stage in *Drosophila melanogaster*"

**SUPPLEMENTARY INFORMATION:**

Table S1.1: Model formulas for each of the response variables. Predictor variables are split into population-level (fixed) effects and group-level (random) effects.

| Response variable | Formula for Population-level effects | Formula for Group-level effects |
| --- | --- | --- |
| Cold Shock Recovery Time | ~ O*M*D + | -1+ O1\|gr(MotherID, by = MD |
| Heat Shock Recovery Time | ~ O*M*D + | -1+ O1\|gr(MotherID, by = MD |
| Survival | ~ O*M*D + | -1+ O1\|gr(MotherID, by = MD |
| Developmental time | ~ O*M*D + | -1+ OT1\|gr(MotherID, by = MD |

Table S1.2: The coefficient of the determination, with 97,5 % credible intervals.

| **Estimate + 95% CI** | **Cold Shock Recovery Time** | **Heat Shock Recovery Time** | **Survival** | **Developmental time** |
| --- | --- | --- | --- | --- |
| **R2** | 0.5278  (0.4636, 0.5822) | 0.6032  (0.5376, 0.6596) | 0.9401  (0. 9345, 0. 9456) | 0.9888  (0. 9886, 0. 9889) |

Table S1.3: Model output for cold shock recovery time, including estimates (Estimate; posterior mean), estimation error (Est.Error; posterior standard deviation), Rhat values for model, and effective sample sizes.

| **Group-Level Effects:** | | | | | | | |
| --- | --- | --- | --- | --- | --- | --- | --- |
| ~MotherID | Number of levels: 175 | | | | | | |
|  | Estimate | Est.Error | l-95% CI | u-95% CI | Rhat | Bulk_ESS | Tail_ESS |
| sd(O_TC:MDC.larva) | 0.32 | 0.07 | 0.19 | 0.46 | 1.00 | 3632 | 5626 |
| sd(O_TH:MDC.larva) | 0.59 | 0.09 | 0.44 | 0.79 | 1.00 | 3219 | 6032 |
| sd(O_TC:MDH.larva) | 0.19 | 0.08 | 0.03 | 0.35 | 1.00 | 1806 | 1289 |
| sd(O_TH:MDH.larva) | 0.38 | 0.09 | 0.23 | 0.57 | 1.00 | 3769 | 4591 |
| sd(O_TC:MDC.adult) | 0.11 | 0.08 | 0.00 | 0.30 | 1.00 | 7097 | 4749 |
| sd(O_TH:MDC.adult) | 0.44 | 0.13 | 0.22 | 0.71 | 1.00 | 4261 | 5062 |
| sd(O_TC:MDH.adult) | 0.10 | 0.08 | 0.00 | 0.30 | 1.00 | 5775 | 4083 |
| sd(O_TH:MDH.adult) | 0.55 | 0.12 | 0.35 | 0.80 | 1.00 | 4701 | 6868 |
| cor(O_TC:MDC.larva,O_TH:MDC.larva) | 0.25 | 0.37 | -0.57 | 0.84 | 1.01 | 406 | 837 |
| cor(O_TC:MDH.larva,O_TH:MDH.larva) | 0.48 | 0.38 | -0.45 | 0.97 | 1.00 | 909 | 935 |
| cor(O_TC:MDC.adult,O_TH:MDC.adult) | -0.00 | 0.58 | -0.95 | 0.95 | 1.00 | 819 | 2283 |
| cor(O_TC:MDH.adult,O_TH:MDH.adult) | 0.00 | 0.58 | -0.95 | 0.96 | 1.02 | 339 | 1127 |
| **Population-Level Effects:** | | | | | | | |
|  | Estimate | Est.Error | l-95% CI | u-95% CI | Rhat | Bulk_ESS | Tail_ESS |
| Intercept | 6.50 | 0.07 | 6.35 | 6.63 | 1.00 | 1884 | 4640 |
| O_TH | 0.08 | 0.12 | -0.16 | 0.33 | 1.00 | 2129 | 3537 |
| M_TH | -0.06 | 0.09 | -0.24 | 0.12 | 1.00 | 2717 | 5221 |
| dev_stageadult | -0.11 | 0.12 | -0.35 | 0.14 | 1.00 | 2666 | 5387 |
| O_TH:M_TH | 0.61 | 0.15 | 0.30 | 0.91 | 1.00 | 2409 | 4317 |
| O_TH:dev_stageadult | 0.13 | 0.20 | -0.26 | 0.52 | 1.00 | 2656 | 4513 |
| M_TH:dev_stageadult | 0.09 | 0.17 | -0.23 | 0.42 | 1.00 | 3315 | 5724 |
| O_TH:M_TH:dev_stageadult | -0.52 | 0.27 | -1.04 | -0.00 | 1.00 | 2705 | 4650 |
| **Family Specific Parameters:** | | | | | | | |
|  | Estimate | Est.Error | l-95% CI | u-95% CI | Rhat | Bulk_ESS | Tail_ESS |
| sigma | 0.44 | 0.02 | 0.41 | 0.48 | 1.00 | 3953 | 6779 |

Table S1.4: Model output for heat shock recovery time, including estimates (Estimate; posterior mean), estimation error (Est.Error; posterior standard deviation), Rhat values for model, and effective sample sizes.

| **Group-Level Effects:** | | | | | | | |
| --- | --- | --- | --- | --- | --- | --- | --- |
| ~MotherID | Number of levels: 144 | | | | | | |
|  | Estimate | Est.Error | l-95% CI | u-95% CI | Rhat | Bulk_ESS | Tail_ESS |
| sd(O_TC:MDC.larva) | 0.43 | 0.11 | 0.24 | 0.69 | 1.00 | 3503 | 5429 |
| sd(O_TH:MDC.larva) | 0.24 | 0.12 | 0.02 | 0.48 | 1.00 | 2674 | 2192 |
| sd(O_TC:MDH.larva) | 0.57 | 0.10 | 0.40 | 0.80 | 1.00 | 3897 | 4789 |
| sd(O_TH:MDH.larva) | 0.30 | 0.13 | 0.04 | 0.55 | 1.00 | 1767 | 1960 |
| sd(O_TC:MDC.adult) | 0.13 | 0.10 | 0.00 | 0.38 | 1.00 | 6428 | 4392 |
| sd(O_TH:MDC.adult) | 0.48 | 0.15 | 0.23 | 0.81 | 1.00 | 3893 | 3674 |
| sd(O_TC:MDH.adult) | 0.15 | 0.11 | 0.01 | 0.41 | 1.00 | 4735 | 3849 |
| sd(O_TH:MDH.adult) | 0.36 | 0.11 | 0.13 | 0.58 | 1.00 | 2706 | 1871 |
| cor(O_TC:MDC.larva,O_TH:MDC.larva) | 0.42 | 0.43 | -0.61 | 0.97 | 1.00 | 3434 | 4806 |
| cor(O_TC:MDH.larva,O_TH:MDH.larva) | -0.02 | 0.33 | -0.70 | 0.62 | 1.00 | 3321 | 3547 |
| cor(O_TC:MDC.adult,O_TH:MDC.adult) | 0.01 | 0.57 | -0.95 | 0.95 | 1.01 | 859 | 2701 |
| cor(O_TC:MDH.adult,O_TH:MDH.adult) | 0.01 | 0.58 | -0.95 | 0.95 | 1.01 | 876 | 2426 |
| **Population-Level Effects:** | | | | | | | |
|  | Estimate | Est.Error | l-95% CI | u-95% CI | Rhat | Bulk_ESS | Tail_ESS |
| Intercept | 6.31 | 0.12 | 6.08 | 6.55 | 1.00 | 2594 | 4437 |
| O_TH | -0.61 | 0.13 | -0.87 | -0.36 | 1.00 | 3235 | 4904 |
| M_TH | -0.16 | 0.16 | -0.49 | 0.16 | 1.00 | 2324 | 4287 |
| dev_stageadult | 0.72 | 0.16 | 0.40 | 1.04 | 1.00 | 3100 | 5078 |
| O_TH:M_TH | -0.02 | 0.19 | -0.40 | 0.35 | 1.00 | 2746 | 4422 |
| O_TH:dev_stageadult | -0.27 | 0.22 | -0.70 | 0.15 | 1.00 | 3642 | 5990 |
| M_TH:dev_stageadult | 0.04 | 0.22 | -0.40 | 0.49 | 1.00 | 2918 | 5017 |
| O_TH:M_TH:dev_stageadult | 0.55 | 0.29 | 0.00 | 1.13 | 1.00 | 3240 | 5445 |
| **Family Specific Parameters:** | | | | | | | |
|  | Estimate | Est.Error | l-95% CI | u-95% CI | Rhat | Bulk_ESS | Tail_ESS |
| sigma | 0.49 | 0.02 | 0.45 | 0.54 | 1.00 | 3575 | 5820 |

Table S1.5: Model output for survival, including estimates (Estimate; posterior mean), estimation error (Est.Error; posterior standard deviation), Rhat values for model, and effective sample sizes.

| **Group-Level Effects:** | | | | | | | |
| --- | --- | --- | --- | --- | --- | --- | --- |
| ~MotherID | Number of levels: 975 | | | | | | |
|  | Estimate | Est.Error | l-95% CI | u-95% CI | Rhat | Bulk_ESS | Tail_ESS |
| sd(O_TC:MDC.egg_to_pupae) | 1.20 | 0.13 | 0.95 | 1.48 | 1.00 | 2959 | 5228 |
| sd(O_TH:MDC.egg_to_pupae) | 0.93 | 0.12 | 0.70 | 1.18 | 1.00 | 4075 | 5761 |
| sd(O_TC:MDH.egg_to_pupae) | 0.99 | 0.14 | 0.74 | 1.27 | 1.00 | 3803 | 6024 |
| sd(O_TH:MDH.egg_to_pupae) | 1.06 | 0.14 | 0.80 | 1.36 | 1.00 | 4647 | 6781 |
| sd(O_TC:MDC.pupae_to_adult) | 1.71 | 0.26 | 1.25 | 2.26 | 1.00 | 3034 | 4782 |
| sd(O_TH:MDC.pupae_to_adult) | 1.93 | 0.27 | 1.44 | 2.50 | 1.00 | 3157 | 5374 |
| sd(O_TC:MDH.pupae_to_adult) | 1.28 | 0.26 | 0.78 | 1.82 | 1.00 | 3006 | 4050 |
| sd(O_TH:MDH.pupae_to_adult) | 1.18 | 0.28 | 0.65 | 1.75 | 1.00 | 2599 | 2849 |
| cor(O_TC:MDC.egg_to_pupae,  O_TH:MDC.egg_to_pupae) | 0.90 | 0.07 | 0.72 | 1.00 | 1.00 | 1434 | 3305 |
| cor(O_TC:MDH.egg_to_pupae,  O_TH:MDH.egg_to_pupae ) | 0.86 | 0.10 | 0.62 | 0.99 | 1.00 | 1395 | 2602 |
| cor(O_TC:MDC.pupae_to_adult,  O_TH:MDC.pupae_to_adult) | -0.17 | 0.22 | -0.59 | 0.27 | 1.00 | 977 | 1603 |
| cor(O_TC:MDH.pupae_to_adult,  O_TH:MDH.pupae_to_adult) | 0.14 | 0.30 | -0.45 | 0.71 | 1.00 | 1575 | 2509 |
| **Population-Level Effects:** | | | | | | | |
|  | Estimate | Est.Error | l-95% CI | u-95% CI | Rhat | Bulk_ESS | Tail_ESS |
| Intercept | 2.36 | 0.13 | 2.11 | 2.63 | 1.00 | 3456 | 4965 |
| O_TH | -0.10 | 0.15 | -0.38 | 0.19 | 1.00 | 4266 | 6084 |
| M_TH | 0.36 | 0.18 | -0.01 | 0.72 | 1.00 | 3561 | 5779 |
| dev_stagepupae_to_adult | 2.38 | 0.33 | 1.76 | 3.04 | 1.00 | 3543 | 4994 |
| O_TH:M_TH | 0.30 | 0.23 | -0.14 | 0.75 | 1.00 | 4132 | 6409 |
| O_TH:dev_stagepupae_to_adult | -0.29 | 0.42 | -1.12 | 0.55 | 1.00 | 3809 | 5350 |
| M_TH:dev_stagepupae_to_adult | -0.81 | 0.42 | -1.61 | 0.02 | 1.00 | 3834 | 5863 |
| O_TH:M_TH:dev_stagepupae_to_adult | -0.09 | 0.54 | -1.15 | 0.98 | 1.00 | 3403 | 6117 |

Table S1.6: Model output for developmental time, including estimates (Estimate; posterior mean), estimation error (Est.Error; posterior standard deviation), Rhat values for model, and effective sample sizes.

| **Group-Level Effects:** | | | | | | | |
| --- | --- | --- | --- | --- | --- | --- | --- |
| ~MotherID | Number of levels: 978 | | | | | | |
|  | Estimate | Est.Error | l-95% CI | u-95% CI | Rhat | Bulk_ESS | Tail_ESS |
| sd(O_TC:MDC.egg_to_pupae) | 0.042 | 0.002 | 0.038 | 0.047 | 1.001 | 3331 | 6248 |
| sd(O_TH:MDC.egg_to_pupae) | 0.030 | 0.002 | 0.026 | 0.034 | 1.001 | 5037 | 6809 |
| sd(O_TC:MDH.egg_to_pupae) | 0.046 | 0.002 | 0.041 | 0.051 | 1.000 | 3176 | 5434 |
| sd(O_TH:MDH.egg_to_pupae) | 0.117 | 0.006 | 0.107 | 0.129 | 1.001 | 2166 | 4370 |
| sd(O_TC:MDC.pupae_to_adult) | 0.046 | 0.003 | 0.041 | 0.051 | 1.002 | 3320 | 5802 |
| sd(O_TH:MDC.pupae_to_adult) | 0.053 | 0.003 | 0.047 | 0.058 | 1.001 | 3037 | 5553 |
| sd(O_TC:MDH.pupae_to_adult) | 0.060 | 0.003 | 0.054 | 0.066 | 1.000 | 2719 | 4594 |
| sd(O_TH:MDH.pupae_to_adult) | 0.153 | 0.007 | 0.139 | 0.168 | 1.000 | 1479 | 2782 |
| cor(O_TC:MDC.egg_to_pupae,  O_TH:MDC.egg_to_pupae) | 0.492 | 0.073 | 0.343 | 0.628 | 1.000 | 3803 | 5608 |
| cor(O_TC:MDH.egg_to_pupae,  O_TH:MDH.egg_to_pupae ) | 0.658 | 0.045 | 0.565 | 0.740 | 1.001 | 1842 | 3520 |
| cor(O_TC:MDC.pupae_to_adult,  O_TH:MDC.pupae_to_adult) | -0.290 | 0.072 | -0.425 | -0.150 | 1.003 | 1773 | 3851 |
| cor(O_TC:MDH.pupae_to_adult,  O_TH:MDH.pupae_to_adult) | 0.462 | 0.058 | 0.344 | 0.574 | 1.010 | 644 | 1330 |
| **Population-Level Effects:** | | | | | | | |
|  | Estimate | Est.Error | l-95% CI | u-95% CI | Rhat | Bulk_ESS | Tail_ESS |
| Intercept | 5.452 | 0.003 | 5.446 | 5.458 | 1.001 | 3021 | 4995 |
| O_TH | -0.921 | -0.921 | -0.927 | -0.915 | 1.001 | 4403 | 7083 |
| M_TH | 0.010 | 0.004 | 0.001 | 0.018 | 1.000 | 2535 | 4373 |
| dev_stagepupae_to_adult | -0.149 | 0.004 | 0.157 | -0.140 | 1.001 | 2619 | 4643 |
| O_TH:M_TH | 0.053 | 0.007 | 0.039 | 0.066 | 1.002 | 2311 | 4474 |
| O_TH:dev_stagepupae_to_adult | -0.007 | 0.006 | 0.019 | 0.005 | 1.001 | 2781 | 4487 |
| M_TH:dev_stagepupae_to_adult | -0.014 | 0.007 | 0.027 | -0.000 | 1.002 | 1966 | 3381 |
| O_TH:M_TH:dev_stagepupae_to_adult | -0.176 | 0.013 | 0.201 | -0.150 | 1.005 | 1169 | 2559 |
| **Family Specific Parameters:** | | | | | | | |
|  | Estimate | Est.Error | l-95% CI | u-95% CI | Rhat | Bulk_ESS | Tail_ESS |
| sigma | 0.052 | 0.000 | 0.052 | 0.053 | 1.000 | 15099 | 7353 |
